# Insights into circovirus host range from the genomic fossil record

**DOI:** 10.1101/246777

**Authors:** Tristan P.W. Dennis, Peter J. Flynn, William Marciel de Souza, Joshua B. Singer, Corrie S. Moreau, Sam J. Wilson, Robert J. Gifford

**Author notes:** These authors contributed equally (Tristan P.W. Dennis, Peter J. Flynn). Corresponding author: Robert J. Gifford MRC-University of Glasgow Centre for Virus Research 464 Bearsden Road Glasgow, Scotland, UK.

## Abstract

A diverse range of DNA sequences derived from circoviruses (family *Circoviridae)* have been identified in samples obtained from humans and domestic animals, often in association with pathological conditions. In the majority of cases, however, little is known about the natural biology of the viruses from which these sequences are derived. Endogenous circoviral elements (CVe) are DNA sequences derived from circoviruses that occur in animal genomes and provide a useful source of information about circovirus-host relationships. In this study we screened genome assemblies of 675 animal species and identified numerous circovirus-related sequences, including the first examples of CVe derived from cycloviruses. We confirmed the presence of these CVe in the germline of the elongate twig ant (*Pseudomyrmex gracilis*), thereby establishing that cycloviruses infect insects. We examined the evolutionary relationships between CVe and contemporary circoviruses, showing that CVe from ants and mites group relatively closely with cycloviruses in phylogenies. Furthermore, the relatively random interspersal of CVe from insect genomes with cyclovirus sequences recovered from vertebrate samples, suggested that contamination might be an important consideration in studies reporting these viruses. Our study demonstrates how endogenous viral sequences can inform metagenomics-based virus discovery. In addition, it raises doubts about the role of cycloviruses as pathogens of humans and other vertebrates.

## Importance

Advances DNA sequencing have dramatically increased the rate at which new viruses are being identified. However, the host species associations of most virus sequences identified in metagenomic samples are difficult to determine. Our analysis indicates that viruses proposed to infect vertebrates (in some cases being linked to human disease) may in fact be restricted to arthropod hosts. The detection of these sequences in vertebrate samples may reflect their widespread presence in the environment as viruses of parasitic arthropods.

## Background

Circoviruses (family *Circoviridae)* are small, non-enveloped viruses with circular, single stranded DNA (ssDNA) genomes ~1.8 to ~2.1 kilobases (kb) in length. Circovirus genomes encode two major proteins: replication-associated protein (Rep) and capsid (Cap), responsible for genome replication and particle formation respectively. Transcription is bidirectional with the *rep* gene being encoded in the forward sense, and the *cap* gene being encoded in the complementary sense (1, 2).

The family *Circoviridae* contains two recognised genera: *Circovirus* and *Cyclovirus* (1). The genus *Circovirus* includes pathogenic viruses of vertebrates, such as porcine circovirus 2 (PCV-2), which causes post-weaning multisystemic wasting syndrome in swine. The genus *Cyclovirus*, by contrast, is comprised entirely of viruses that have been identified only via sequencing, and for which host species associations are less clear. Nevertheless, cycloviruses have frequently been associated with pathogenic conditions in humans and domestic mammals. For example, cyclovirus sequences have been detected in the cerebrospinal fluid of humans suffering from acute central nervous system disease in Vietnam and Malawi (3, 4). Cyclovirus sequences have also been reported in association with numerous other outbreaks of disease in humans and domestic mammals (5–7).

Sequences derived from circoviruses have been shown to be present in the genomes of many eukaryotic species (8, 9). These ‘endogenous circoviral elements’ (CVe) are thought to be derived from the genomes of ancient circoviruses that were – by one means or another - ancestrally integrated into the nuclear genome of germline cells (10). CVe can provide unique information about the long-term co-evolutionary relationships between viruses and hosts - for example the identification of ancient CVe in vertebrate genomes shows that viruses in the genus *Circovirus* have been co-evolving with vertebrate hosts for millions of years (11).

We recently reported the results of a study in which we systematically screened vertebrate whole genome sequence (WGS) data for CVe (11). Here, we expanded this screen to include a total 675 animal genomes, including 307 invertebrate species. Via screening, we identified novel examples of sequences derived from circoviruses, cycloviruses, and the more divergent CRESS-DNA group. We examine the phylogenetic relationships between these sequences, well-studied circovirus isolates, and circovirus-related sequences recovered via metagenomic sequencing of environmental samples or animal tissues. Our analysis raises important questions about the origins of cyclovirus sequences in samples derived from humans and other mammals, and their role in causing disease in these hosts.

## Materials and Methods

### Sequence data

Whole genome sequence (WGS) assemblies of 675 species (**Table S1**) were downloaded from the National Center for Biotechnology Information (NCBI) website. We obtained a representative set of sequences for the genus *Circovirus* and a non-redundant set of vertebrate CVe sequences from an openly accessible dataset we compiled in our previous work (11). This dataset was expanded to include a broader range of sequences in the family *Circoviridae*, including representative species in the *Cyclovirus* genus, and the more distantly related CRESS-DNA viruses (**Table S2**). We used GLUE - an open, data-centric software environment specialized in capturing and processing virus genome sequence datasets (12) - to collate the sequences, alignments and associated data used in this investigation. These data are available in a publicly accessible online repository: https://github.com/giffordlabcvr/DIGS-for-EVEs.

### Genome screening in silico

Genome screening *in silico* was performed using the database-integrated genome screening (DIGS) tool (13). The DIGS procedure comprises two steps. In the first, the basic local alignment search tool (BLAST) program (14) is used to search a genome assembly file for similar to a particular ‘probe’ (i.e. a circovirus Rep or Cap polypeptide sequence). In the second, sequences that produce statistically significant matches to the probe are extracted and classified by BLAST-based comparison to a set of virus reference genomes (see **Table S2**). Results are captured in a MySQL database.

Newly identified CVe identified in this study were assigned a unique identifier (ID), following a convention we established previously (11). The first component is of the ID the classifier ‘CVe’. The second is a composite of two distinct subcomponents separated by a period: the name of CVe group (usually derived from the host group in which the element occurs in (e.g. Carnivora), and the second is a numeric ID that uniquely identifies the insertion. Orthologous copies in different species are given the same number, but are differentiated using the third component of the ID that uniquely identifies the species from which the sequence was obtained. Unique numeric IDs were assigned to novel CVe with reference to those used in the previously assembled dataset (11).

### Alignments and phylogenetic analysis

Multiple sequence alignments were constructed using MUSCLE (15), RevTrans 2.0 (16), MACSE (17) and PAL2NAL (18). Manual inspection and adjustment of alignments was performed in Se-Al (19) and AliView (20). Phylogenies were reconstructed using maximum likelihood as implemented in IQ-TREE (21) and the VT+ G4 protein substitution model (22) as selected using ProTest (23), with support assessed using 1000 nonparametric bootstrap replicates.

### Amplification and sequencing

Genomic DNA was extracted from ant tissue samples following the Moreau protocol (24) and a DNAeasy Blood & Tissue Kit (Qiagen). PCR amplification of CVe-*Pseudomyrmex* was performed using two sets of primer pairs designed with Primer3 (http://bioinfo.ut.ee/primer3–0.4.0/), each comprising one primer anchored in the CVe sequence, and another anchored in the genomic flanking sequence. Primer Pair 1 amplified a sequence that was 694 bp long and Primer Pair 2 amplified a sequence that was 286bp long. Primers were tested using illustra PuReTaq Ready-To-Go PCR Beads (GE Lifesciences). A temperature gradient PCR was performed to assess the optimum annealing temperature for the specific primer pairs. PCR was then performed using the genomic DNA ant extractions. The PCR conditions for this run were: an initial denature stage of 5 minutes at 95°C, 30 cycles of 30 seconds denaturing at 95°C, 30 seconds annealing at 49.7°C for Primer Pair 1 and 62°C for Primer Pair 2, and an extension at 72°C for 1 minute, then after 30 cycles a final extension at 72°C for 5 minutes. Each run included a negative control. Amplification products (800–1000bp) for each PCR reaction were excised and run on agarose gels. Bands of the expected size were excised, purified and sequenced via Sanger sequencing (25).

## Results

### Identification of CVe in animal genomes

We screened WGS data of 675 animal species (**Table S1**) *in silico* to identify sequences related to circoviruses. We identified 300 circovirus-related sequences in total, 76 of which have not been reported previously (Table 1, **Table S3**). To investigate the novel sequences identified in our screen, each was virtually translated and incorporated into a multiple sequence alignment that included a representative set of previously reported circoviruses and CVe (**Table S2**). Incorporation of CVe sequences into an alignment provided a basis for determining their genetic structure and investigating their phylogenetic relationships to circoviruses.

**Table 1.** Novel CVe identified in this study.

| Common name | Scientific name | Class | Order | # Seqs<sup>a</sup> | Status<sup>b</sup> | Intact<sup>c</sup> |
| --- | --- | --- | --- | --- | --- | --- |
| **Circovirus** |  |  |  |  |  |  |
| Tomato clownfish | *Amphiprion frenatus* | **Vertebrata** | Perciformes | 1 | CVe | No |
| Elephantfish | *Paramormyrops kingsleyae* | **Vertebrata** | Osteoglossiformes | 1 | CVe | No |
| **Cyclovirus** |  |  |  |  |  |  |
| Asian bee mite | *Tropilaelaps mercedesae* | **Arthropoda** | Arachnida | 7 | CVe | No |
| Varroa mite | *Varroa destructor* | **Arthropoda** | Arachnida | 19 | CVe | **Yes** |
| Elongate twig ant | *Pseudomyrmex gracilis* | **Arthropoda** | Insecta | 1 | CVe | **Yes** |
| **CRESS** |  |  |  |  |  |  |
| Myxosporean parasite | *Thelohanellus kitauei* | **Cnidaria** | Myxosporea | 1 | CVe | **Yes** |
| Philippine horse mussel | *Modiolus philippinarum* | **Mollusca** | Bivalvia | 4 | CVe | **Yes** |
| Mediterranean mussel | *Mytilus galloprovincialis* | **Mollusca** | Bivalvia | 4 |  | Yes |
| Freshwater snail | *Biomphalaria glabrata* | **Mollusca** | Gastropoda | 1 |  | Yes |
| Tribble's cone | *Conus tribblei* | **Mollusca** | Gastropoda | 3 |  | Yes |
| Western predatory mite | *Galendromus occidentalis* | **Arthropoda** | Arachnida | 1 |  | Yes |
| Phytoseiid predatory mite | *Metaseiulus occidentalis* | **Arthropoda** | Arachnida | 1 |  | Yes |
| Brown recluse spider | *Loxosceles reclusa* | **Arthropoda** | Arachnida | 19 | CVe | **Yes** |
| Scarab beetle | *Oryctes borbonicus* | **Arthropoda** | Insecta | 1 | CVe | **Yes** |
| Drifting brine fly | *Ephydra gracilis* | **Arthropoda** | Insecta | 10 |  | Yes |
| Alkali fly | *Ephydra hians* | **Arthropoda** | Insecta | 8 |  | Yes |
| Amphipod crustacean | *Parhyale hawaiiensis* | **Arthropoda** | Malacostraca | 3 | CVe | No |
| Sea louse | *Caligus rogercresseyi* | **Arthropoda** | Maxillopoda | 4 |  | Yes |
| Tadpole shrimp | *Triops cancriformis* | **Arthropoda** | Branchiopoda | 1 |  | Yes |
| Pork tapeworm | *Taenia solium* | **Cestoda** | Cyclophyllidea | 3 | CVe | **Yes** |

**Footnote:** <sup>a</sup> Number of distinct sequences disclosing similarity to circovirus proteins were identified in species WGS data. <sup>b</sup> Species genomes that were confirmed as containing CVe are indicated, based on the presence in WGS assemblies of at least one contig containing regions of circovirus homology flanked by >3Kb of genomic sequence. <sup>c</sup> Column indicates which species contained circovirus-derived sequences in which protein-coding potential was maintained across the entire length of the detected region of circovirus homology, and this region was at least >200nt in length.

All of the newly identified sequences were derived from *rep* – no novel sequences derived from circovirus *cap* genes were detected. We identified two novel CVe derived from viruses in the genus *Circovirus* in fish genomes (Table 1). One of these, identified in the tomato clownfish (*Amphiprion frenatus*), appeared to an ortholog of a CVe locus previously identified in other perciform fish (11). The other, identified in a mormyrid fish, was clearly related to CVe previously identified in ray-finned fish (11, 26), but as it comprised a relatively short fragment of the *rep* gene, its more precise phylogenetic relationship to these CVe could not be determined with confidence.

We identified 93 circovirus-related sequences in invertebrate genome assemblies, 71 of which have not been reported previously (Table 1, **Table S3**). Of these, a relatively high proportion exhibited coding potential. Some occurred on short contigs, and could potentially have been derived from contaminating virus. However, we found that in many cases, at least one of the circovirus-related sequences identified in a WGS assembly was incorporated into a contig that was easily large enough to contain an entire circovirus genome, and flanked by >3 kilobases (Kb) of genomic sequence, and thus very likely to represent CVe. On this basis, we estimate that 60 of the 93 sequences we identified in invertebrate genomes are likely to be derived from CVe. Sequences that occurred on short contigs - particularly those that lacked any in-frame stop codons or frameshifts (see Table 1) - might instead be derived from contaminating virus.

Maximum likelihood (ML) phylogenies were reconstructed using an alignment of Rep proteins, and disclosed two robustly supported, monophyletic clades corresponding to the *Circovirus* and *Cyclovirus* genera (1). In line with our previous investigations (11), we found that all Rep-related sequences from vertebrate WGS grouped with circoviruses, with the exception of a highly divergent sequence identified in the genome of a jawless vertebrate (the hagfish *Eptatretus burgeri)*. All sequences derived from invertebrate WGS grouped with cycloviruses, or with divergent CRESS-DNA viruses (e.g. Avon-Heathcote Estuary associated circular virus 24 (27)). CRESS-DNA virus-like sequences from distinct species tended to emerge on relatively long branches, and bootstrap support for branching patterns in this region of the phylogeny were generally quite low. The low resolution in this part of the phylogeny likely reflects the lack of adequate sampling of viruses from invertebrate species.

Some sequences from invertebrate WGS were observed to cluster with cycloviruses in phylogenies. Some of these sequences occur within relatively short contigs. However, others occur on relatively large contigs (e.g. > 1 kilobase) that were easily large enough contain CVe and genomic flanking sequences, but similarity to previously sequenced arthropod genomes could not be detected. They include CVe identified in WGS data of two parasitic mite species: *Varroa destructor* and *Tropilaelaps mercedesae*, and in the genome of the elongate twig ant *(Pseudomyrmex gracilis)*. This latter element – hereafter referred to as CVe-*Pseudomyrmex* - was no more distantly related to contemporary cycloviruses than many of them are to one another, including some that are associated with vertebrates (at least superficially) (Figure 1). Because this seemed a little surprising, we sought to confirm the presence of CVe-*Pseudomyrmex* in the twig ant germline. We obtained genomic DNA from four species of ant within the *Pseudomyrmex* genus *(P. gracilis, P. elongatus, P. spinicola*, and *P. oculatus)*, including three distinct populations of *P. gracilis*, We then used polymerase chain reaction (PCR) to amplify a region encompassing part of the CVe, and part of the genomic flanking sequence. We obtained an amplicon of the expected size in all three DNA samples of *P.gracilis*, all other samples were negative (Figure 2). Sequencing of the amplicon confirmed that it was derived the genomic locus containing the CVe, and contained both a portion of the CVe and a region of genomic flanking sequence.

**Figure 1.**
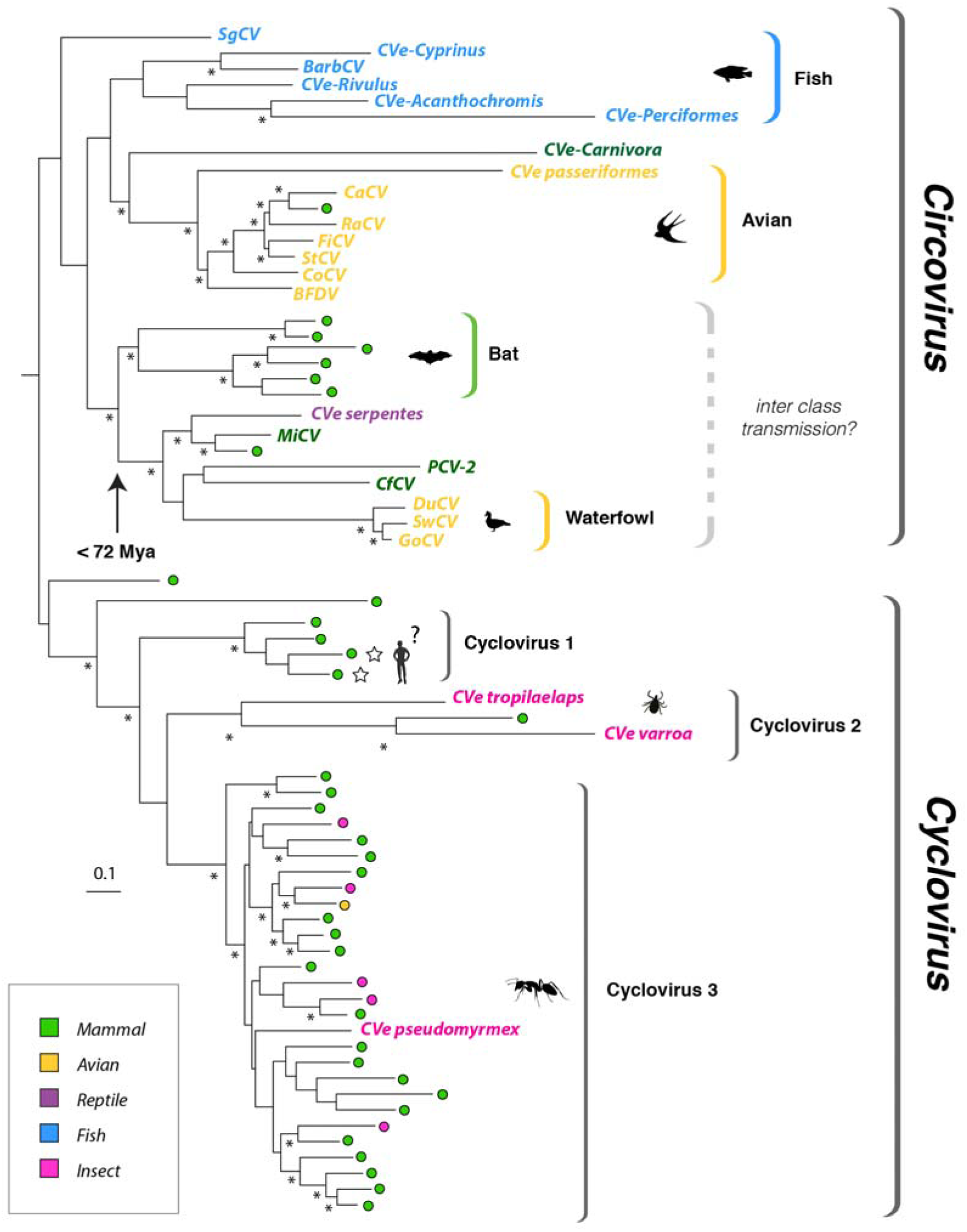
Phylogeny of exogenous and endogenous circovirus Rep sequences. Maximum-likelihood phylogeny reconstructed from an alignment of replication-associated protein (Rep) sequences. The tree is mid-point rooted, asterisks indicate nodes with >70% bootstrap support. Scale bar indicates evolutionary distance in substitutions per site. Sequences derived from metagenomic samples are indicated by colored circles. Taxa names are shown for sequences derived from viruses and CVe. All taxa are coloured to indicate associations with host species groups, as shown in the key. Stars indicate viral taxa that have been linked to human disease. See **Figure S1** for accession numbers of all taxa shown here. The arrow indicates an age calibration inferred for a clade within the *Circovirus* genus. Mya=million years ago.

**Figure 2.**
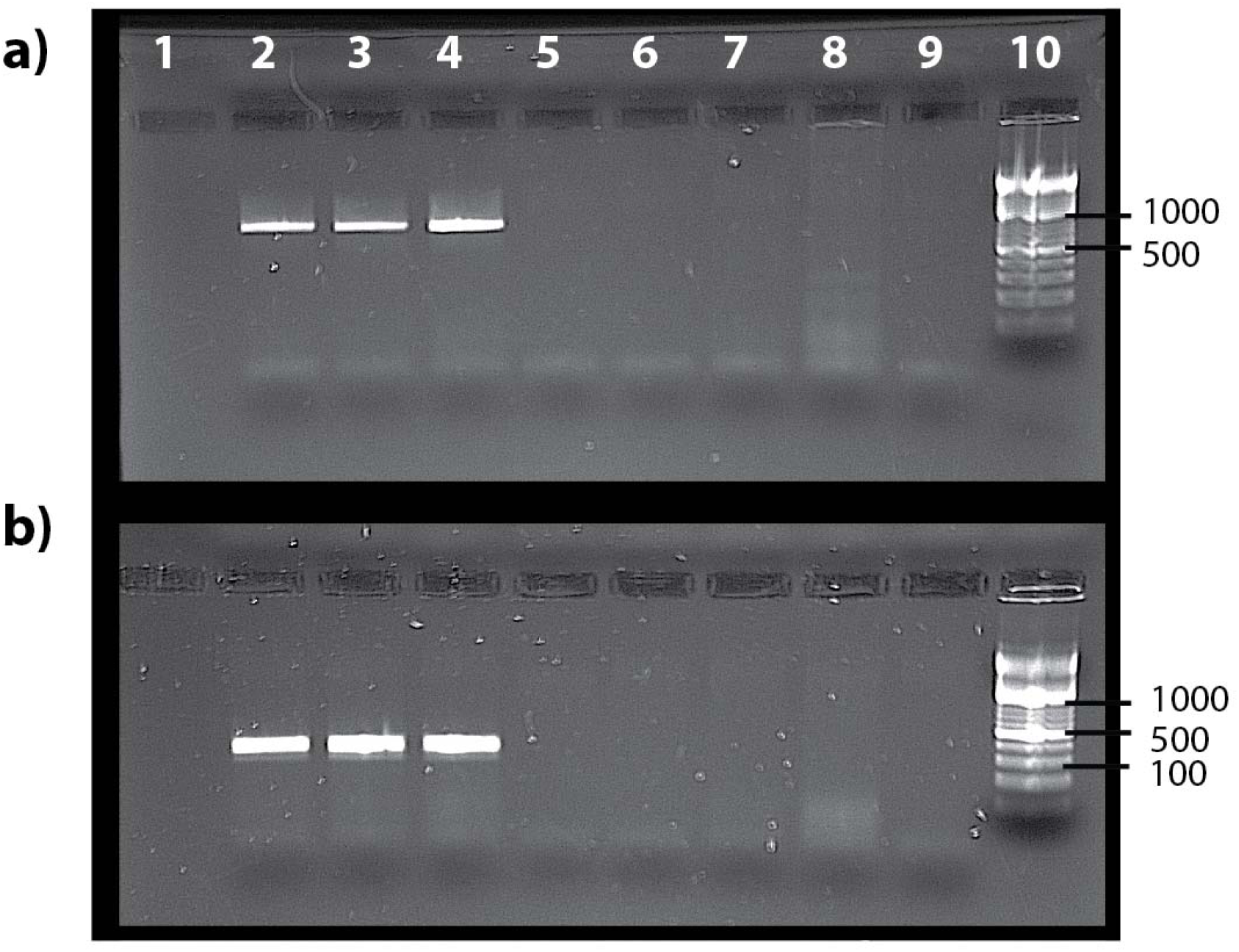
PCR confirmation of *CVe-Pseudomyrmex* presence in three populations of *Pseudomyrmex gracilis*. Panel (a) shows the results of amplification using primer pair 1 (694bp product), panel (a) shows the results of amplification using primer pair 2 (286bp product). Columns: (1) negative control; (2) *Pseudomyrmex gracilis*, from the Florida Keys; (3) *P. gracilis* from mainland Florida, USA; (4) *P. gracilis* from Texas, USA; (5) *P. elongatus* from the Florida Keys, USA; (6) *P. spinicola* from Guanacaste Province, Costa Rica; (7) *P. oculatus* from Cusco, Peru; (8) *Cephalotes atratus*, from Cusco Peru; (9) negative control; (10) ladder.

### Mapping the host associations of circoviruses and cycloviruses

In phylogenies based on Rep, clades corresponding to the *Circovirus* and *Cyclovirus* genera contained a mixture of; (i) CVe from WGS assemblies; (ii) sequences obtained from virus isolates; (iii) sequences obtained from metagenomic samples (Figure 1, **Table S1**). Among circoviruses, associations at the level of class appear relatively stable. For example, beak and feather disease virus (BFDV) groups robustly with a CVe that entered the germline of passeriforme birds ~38 Mya (11), while barbel circovirus groups (BarbCV) groups robustly with CVe from the genome of the golden line barbell, in a well-supported clade containing numerous CVe from ray-finned fish. The only sequence that superficially seems to contradict this pattern is ‘chimpanzee circovirus’, which groups robustly with avian viruses. However, the name of this sequence is misleading – although it was recovered from chimpanzee feces but no host association is known – indeed, the possibility that it might derive from an avian circovirus was noted at the time it was reported (28).

We observed three well-supported sublineages in the clade corresponding to the *Cyclovirus* genus, here labelled cyclovirus 1–3 (see Figure 1). For cycloviruses, the only confirmed host associations come from the *CVe-Pseudomyrmex* sequence reported above. Many of the cyclovirus sequences that have been identified via metagenomic sequencing are associated with arthropod species, such as dragonflies (29). However, others are associated with vertebrates, having names such as ‘bat cyclovirus’. Cyclovirus 1 is exclusively comprised of viruses from vertebrate samples. In cyclovirus groups 2 and 3, however, sequences from vertebrate and invertebrate samples are extensively intermingled (Figure 1), and clade structure does not reflect these host associations in any obvious way. Sequences from each host group appear to be dispersed randomly, and the branch lengths separating vertebrate from invertebrate viruses (and CVe) are relatively short in many cases.

## Discussion

In this study, we screened *in silico* 675 animal genomes and identified numerous sequences related to circoviruses, including many that have not been reported previously. We examine the phylogenetic relationships between these sequences, well-studied circovirus isolates, and circovirus sequences recovered via metagenomic sequencing of environmental samples or animal tissues.

Most of the novel circovirus sequences reported here were identified in invertebrate genome assemblies. Many were highly divergent, and are likely derived from uncharacterised CRESS-DNA virus lineages that infect invertebrate species. All of the newly identified sequences identified in our study were derived from *rep* genes – we did not detect any novel CVe disclosing homology to circovirus *cap* genes. The preponderance of CVe derived from *rep* versus those derived from *cap* might reflect the greater heterogeneity of capsid sequences in general, which might lead to these sequence being generally harder to detect – certainly, in the case of the more divergent invertebrate viruses, it is possible that the *cap* genes found in some lineages might not share any sequence homology with those sequenced previously. However, we note that even among CVe derived from viruses in the genus *Circovirus*, within which capsid sequences are comparatively conserved, sequences derived from *rep* are around twice as common as those derived from *cap*.

Among the factors that may have influenced the structure and types of CVe that we observe in animal germlines are selection pressures that have led to these sequences being co-opted or exapted by host species. Interestingly, several of the confirmed CVe in our study lacked frameshifting mutations or in-frame stop codons (see Table 1), indicating that they have been evolving under purifying selection relatively recently. We confirmed the presence of one such CVe (CVe-*Pseudomyrmex*) in three populations of *Pseudomyrmex gracilis* (Figure 2). The fact that we did not detect the *CVe-Pseudomyrmex* sequence in other members of the genus suggests it was incorporated into the *P. gracilis* germline after this species diverged from *P. elongatus, P. spinicola, and P. oculatus* in the Miocene epoch (30). However, we cannot completely rule out that *Pseudomyrmex* is actually older, since the failure to obtain an amplicon in other *Pseudomyrme*x species could be accounted for in other ways (e.g. sequence divergence in the regions targeted by PCR primers). Nevertheless, the occurrence of an apparently fixed, intact, and expressed circovirus *rep* gene in an ant genome provides further evidence that these genes have been co-opted or exapted by host species for as yet unknown functions. Functional genomic studies in insects indicate that endogenous viral element (EVE) sequences have been co-opted into RNA-based systems of antiviral immunity (31). Thus, one possible explanation accounting for the conservation of this sequence in CVe-*Pseudomyrmex* is that it is involved in immune defence, although this would not necessarily require maintenance of an intact coding sequence.

Our study allowed the host associations of circoviruses and CVe to be examined in the context of their evolutionary relationships. With respect to this, the grouping of sequences for which the host associations are well established (i.e. CVe and viruses that have been investigated using methods besides only sequencing) relative to sequences recovered from metagenomic samples, was revealing. Prior to this study, the only host associations that had been robustly demonstrated were within the genus *Circovirus*. Circoviruses have been isolated from vertebrates, and in phylogenies based on Rep proteins, these isolates group together with vertebrate CVe in a well-supported clade. Furthermore, the host associations of circoviruses appear quite stable, with ancient CVe from particular host groups (e.g. orders, classes) sometimes seen grouping together with contemporary viruses from the same host groups (Figure 1). Within the *Circovirus* clade there is only one sequence that seems to contradict this pattern. This sequence was recovered from a stool sample and thus - as was observed in when it was first reported (28) – is likely to reflect environmental contamination.

The limited evidence available regarding the zoonotic potential of circoviruses suggests they lack the capacity to be transmitted between relatively distantly related hosts (i.e. hosts in distinct classes or orders). For example, during the 1990s and early 2000s, porcine circovirus 1 (PCV-1) was inadvertently introduced into batches of live attenuated rotavirus vaccine as an adventitious agent. These vaccines were administered to millions of people (32), yet PCV-1 is not thought to have infected any humans as a result, indicating that powerful barriers to cross-species transmission are probably in effect.

Nevertheless, recent studies have identified some surprising cases wherein phylogenetic trees indicate apparent transmission of viruses between vertebrate classes (33). We see evidence for potential inter-class transmission within one *Circovirus* subclade that contains sequences obtained from waterfowl and mammals (including mink, bats, dogs and pigs) as well as CVe from reptile genomes (see Figure 1). Within this subclade, viruses of mink are robustly separated from porcine, canine and waterfowl viruses by a CVe that was incorporated into the serpentine germline >72 million years ago (11). Thus, the phylogeny indicates that at least one interclass transmission event is likely to have occurred within this clade. However, it should be noted that while the clustering patterns observed in this clade do suggest potential transmission of circoviruses between vertebrate classes, they also indicate that such events have occurred relatively infrequently during evolution, since the CVe in snake genomes provides a minimum for the entire clade (assuming the root of this clade is as depicted in Figure 1). Moreover, since clustering patterns that superficially appear to indicate host switches can be accounted for by multiple alternative evolutionary scenarios (34, 35), caution is advisable when using phylogenetic approaches to infer the relationships between parasites and their hosts, especially when sampling is limited.

If cross-species transmission of circoviruses between distinct mammalian orders does not occur readily, then transmission between arthropod and vertebrate hosts appears extremely unlikely. However, if we take the reported host species associations of cycloviruses at face value, we might conclude it occurs frequently, particularly within the ‘cyclovirus 3’ lineage (Figure 1). Importantly, CVe*-Pseudomyrmex* groups robustly within this lineage, and since all other taxa within the *Cyclovirus* clade have been recovered via metagenomic screening, this provides the first unambiguous evidence of a host-association for cycloviruses, establishing with a high degree of confidence that they do indeed infect arthropods, or at the very least, have done so in the past. Furthermore, since the only proven associations for cycloviruses so far are with arthropods, contamination of vertebrate samples with viruses derived from arthropods is perhaps the most parsimonious explanation for the host associations observed here. Contamination from arthropod sources such as dust mites can presumably occur fairly easily given their ubiquity (36). Intriguingly with respect to this, we identified putative cyclovirus CVe in the genomes of two distinct mite species (Figure 1, Table 1).

Since there is always a risk of being misled by contamination when identifying viruses via sequencing-based approaches, we propose that host associations of circoviruses identified via sequencing should be viewed with caution where they are found to strongly contradict established host associations within well-defined clades, particularly at higher taxonomic levels (e.g. phylum, class, order). Whereas the weight of evidence may favour cyclovirus clades 2 and 3 being exclusively arthropod viruses that frequently contaminate vertebrate samples, the status of the cyclovirus 1 lineage is perhaps more equivocal. This basal lineage is comprised exclusively of sequences obtained from mammalian samples, and includes cycloviruses proposed to cause disease in humans (cyclovirus VN and human cyclovirus VS5700009). Conceivably, these sequences could represent a mammal or vertebrate-specific lineage of cycloviruses that is distinct from arthropod-infecting lineages. Notably, however false-positive detection of human cyclovirus VS5700009 has been reported (37).

Virus sequences recovered from metagenomic samples can be investigated by examining their phylogenetic relationships to other viruses for which host associations have been established. The work performed here demonstrates the utility of endogenous virus sequences in this process. This approach can be generalized to inform metagenomics-based virus discovery and diversity mapping efforts for any virus group that has generated endogenous sequences.

## ACKNOWLEDGEMENTS

TPWD, JBS, SJW, and RJG were supported by a grant from the UK Medical Research Council (No. MC_UU_12014/10). WMS was supported by the Fundação de Amparo à Pesquisa do Estado de São Paulo, Brazil (Scholarships No. 12/24150–9; 15/05778–5; 17/13981–0). CSM and PJF were supported by a grant from the National Science Foundation (NSF DEB 1442316).

